# Detection and functional resolution of soluble multimeric immune complexes by a comprehensive FcγR reporter cell panel

**DOI:** 10.1101/2020.11.11.378232

**Authors:** Haizhang Chen, Andrea Maul-Pavicic, Martin Holzer, Magdalena Huber, Ulrich Salzer, Nina Chevalier, Reinhard E. Voll, Hartmut Hengel, Philipp Kolb

## Abstract

Fc-gamma receptor (FcγR) activation by soluble IgG immune complexes (sICs) represents a major mechanism of inflammation in certain autoimmune diseases such as systemic lupus erythematosus (SLE). A robust and scalable test system allowing for the detection and quantification of sIC bioactivity is missing. Previously described FcγR interaction assays are limited to certain FcγRs, lack scalability and flexibility, are not indicative of receptor activation or lack sensitivity towards sIC size. We developed a comprehensive reporter cell panel detecting individual activation of FcγRs from humans and the mouse. The reporter cell lines were integrated into an assay format that provides flexible read-outs enabling the quantification of sIC reactivity via ELISA or a fast detection using flow cytometry. This identified FcγRIIA(H) and FcγRIIIA as the most sIC-sensitive FcγRs in our test system. Applying the assay we demonstrate that sICs versus immobilized ICs are fundamentally different FcγR-ligands with regard to FcγR preference and signal strength. Reaching a detection limit in the very low nanomolar range, the assay proved also to be sensitive to sIC stoichiometry and size enabling for the first time a complete reproduction of the Heidelberger-Kendall precipitation curve in terms of immune receptor activation. Analyzing sera from SLE patients and mouse models of lupus and arthritis proved that sIC-dependent FcγR activation has predictive capabilities regarding severity of SLE disease. The new methodology provides a sensitive, scalable and comprehensive tool to evaluate the size, amount and bioactivity of sICs in all settings.

**One Sentence Summary:** In this study we established a comprehensive FcγR reporter cell assay enabling the detection and quantification of soluble immune complexes generated in experimental and clinical settings.

## Introduction

Immunoglobulin G (IgG) is the dominant immunoglobulin isotype in chronic infections and in various antibody-mediated autoimmune diseases. The multi-faceted effects of the IgG molecule rely both on the F(ab) regions, which recognize a specific antigenic determinant to form immune complexes (ICs), and the constant Fc region (Fcγ), which is detected by effector molecules like the Fcγ receptors (FcγRs) found on most cells of the immune system or C1q of the complement system. When IgG binds to its antigen ICs are formed, which, depending on the respective antigen, are either matrix/cell-bound or soluble (sICs). The composition of sICs is dependent on the number of epitopes recognized by IgG on a single antigen molecule and the ability of the antigen to form multimers. Fcγ-FcγR binding is necessary but not sufficient to activate FcγRs since physical receptor cross-linking is required for receptor triggering (Duchemin et al, 1994; Luo et al, 2010; Patel et al, 2019). IgG opsonized infected cells or microorganisms are readily able to cross-link FcγRs (Bruhns et al, 2009; Lux et al, 2013). This initiates various signalling pathways (Greenberg et al, 1994; Kiefer et al, 1998; Luo et al, 2010) which in turn regulate immune cell effector functions (Bournazos et al, 2017; Nimmerjahn & Ravetch, 2010). It is also suggested that sICs can dynamically tune FcγR triggering, implying that changes in sIC size directly impact strength and duration of FcγR responses (Lux et al, 2013). However, the molecular requirements are largely unknown and a translation to bioactivity of the paradigmatic Heidelberger-Kendall precipitation curve, describing that sIC size depends on the antigen:antibody ratio, is so far missing (Heidelberger & Kendall, 1929; Heidelberger & Kendall, 1935).

Among all type I FcγRs, FcγRIIB (CD32B) is the only inhibitory receptor, signalling via immunoreceptor tyrosine-based inhibitory motifs (ITIMs), while the activating receptors are associated with immunoreceptor tyrosine-based activation motifs (ITAMs). Another exception is FcγRIIIB (CD16B), which is glycosylphosphatidylinositol (GPI)-anchored and lacks a signalling motif (Bruhns, 2012; Bruhns & Jonsson, 2015; Nimmerjahn & Ravetch, 2006; Nimmerjahn & Ravetch, 2008). Still, FcγRIIIB is widely accepted to be a neutrophil activating receptor, e.g. by cooperating with other FcγRs such as FcγRIIA (Vossebeld et al, 1997). FcγRI (CD64) is the only high affinity FcγR binding also to monomeric IgG, while all other FcγRs only efficiently bind to complexed, i.e. antigen-bound IgG (Bruhns, 2012; Bruhns & Jonsson, 2015; Lu et al, 2018). Activation of FcγRs leads to a variety of cellular effector functions elicited by several immune cells such as natural killer (NK) cells via FcγRIIC/FcγRIIIA, monocyte-derived cells via FcγRI/FcγRIIB/FcγRIIIA, granulocytes via FcγRI/FcγRIIA/FcγRIIIB, platelets via FcγRIIA and B cells via FcγRIIB. Consequently, FcγRs connect and regulate both the innate and adaptive branches of the immune system. Various factors have been shown to influence IC-dependent FcγR activation profiles, including FcγR-Fcγ binding affinity and avidity (Koenderman, 2019), IgG subclass, glycosylation patterns and genetic polymorphism (Bruhns et al, 2009; Pincetic et al, 2014; Plomp et al, 2017; Vidarsson et al, 2014), stoichiometry of antigen-antibody-ratio (Berger et al, 1996; Lux et al, 2013; Pierson et al, 2007) and FcγR clustering patterns (Patel et al, 2019). Specifically, Asn297-linked glycosylation patterns of the IgG Fc domain initiate either pro- or anti-inflammatory effector pathways by tuning the binding affinity to activating versus inhibitory FcγRs, respectively (Bohm et al, 2014). However, despite being explored in proof-of-concept studies, the functional consequences of these ligand features on a given FcγR are still not fully understood. Therefore, there is an obvious need for an assay platform allowing for the systematic assessment of IC-mediated FcγR activation.

sICs and immobilized ICs represent unequal and discrete stimuli for the immune system (Fossati et al, 2002; Granger et al, 2019). Soluble circulating ICs are commonly associated with certain chronic viral or bacterial infections (Wang & Ravetch, 2015; Yamada et al, 2015) and autoimmune diseases, such as systemic lupus erythematosus (SLE) or rheumatoid arthritis (RA) (Antes et al, 1991; Koffler et al, 1971; Zubler et al, 1976). When deposited and accumulating in tissues, sICs can cause local damage due to inflammatory responses, classified as type III hypersensitivity (Rajan, 2003). Compared with immobilized local ICs, which recruit immune cells causing tissue damage (Mayadas et al, 2009; Mulligan et al, 1991; Ward et al, 2016), sIC related disorders are characterized by systemic inflammation which is reflected by immune cell exhaustion and senescence (Bano et al, 2019; Chauhan, 2017; Tahir et al, 2015). In order to resolve sIC-dependent activation of FcγRs in greater detail, we developed a scalable reporter system suited for two high throughput readouts, capable of quantifying and distinguishing the activation of single FcγRs. As the assay is also sensitive to stoichiometry and sIC size, we were able to translate the Heidelberger-Kendall precipitation curve to FcγR bioactivity. Compared to currently available ELISA assays detecting sICs by their affinity to C1q-CIC (circulating immune complexes) or C3d, the assay system presented below is strictly specific for IgG immune complexes and integrates sICs of all sizes into single Fcγ receptor bioactivity. Applying reporter cell lines enables very high sensitivity in the low nanomolar range, as signals are biologically amplified compared to biochemical binding based read-outs. Finally, we applied the assay to a clinical setting, measuring sICs in sera from SLE patients. A reporter cell panel expressing murine FcγRs revealed the detection of sICs in the serum of autoimmune-prone diseased mice in preclinical models of lupus and arthritis. Prospectively, this methodology could be instrumental as an experimental and clinical toolbox to unveil sIC-mediated FcγR activation in various autoimmune or infectious diseases.

## Results

### Experimental assay setup

The assay used in this study was adapted from a previously described cell-based FcγR activation test system designed to measure receptor activation in response to opsonized virus infected cells (Corrales-Aguilar et al, 2013; Kolb et al, 2021) and therapeutic Fc-fusion proteins (Lagasse et al, 2019). We refined the assay to enable selective detection of sICs and expanded the reporter cell line-up (FcγRI: Acc# LT744984; FcγRIIA (131R): Acc# M28697; FcγRIIA(131H): Acc# XP_011507593; FcγRIIB/C: Acc# LT737639; FcγRIIIA(176V): Acc# LT737365; FcγRIIIB(176V): Acc# O75015). Ectodomains of FcγRIIB and FcγRIIC are identical. Second generation reporter cells were generated to improve stable expression of chimeric FcγRs compared to the transfectants used in the original assay (Corrales-Aguilar et al, 2013). To this end, mouse BW5147 cells were transduced as described previously via lentiviral transduction (Corrales-Aguilar et al, 2013; Halenius et al, 2011; Van den Hoecke et al, 2017). Human FcγR expression on transduced cells after puromycin selection and two consecutive cell sorting steps was assessed by flow cytometry (Fig. 1A). FcγR activation is measured by surface CD69 expression after 4h of incubation using high thoughput flow cytometry or by quantification of IL-2 secretion after 16 h of incubation using ELISA. Suspension of IgG or sICs in the liquid phase is enforced by pre-incubation of a 96 well ELISA microtiter plate with PBS/FCS blocking buffer (Fig. 1B). To this end, we compared graded concentrations of FCS in the blocking reagent and measured the threshold at which IgG was no longer adsorbed to the plate and stayed abundantly in solution. FCS supplementation to 1% (v/v) or higher is sufficient to keep IgG antibodies in solution. We then set out to test if immobilized IgG can be used as an operational surrogate for IgG-opsonized cells or immobilized ICs with regard to FcγR activation as suggested previously (Tanaka et al, 2009). We found no qualitative difference in FcγR activation between immobilized Rtx, immobilized ICs (Rtx + rec. CD20) or Rtx-opsonized 293T-CD20 cells (Fig. 1C). In contrast, sICs formed by monomeric CD20 antigen (aa 141-188) and Rtx failed to activate FcγRs even at very high ligand concentrations. We concluded that FcγR-crosslinking by sICs is only achieved by multivalent antigens but not dimeric ICs. Of particular note, to reliably and accurately differentiate between strictly soluble and immobilized including aggregated triggers using this assay, reagents for the generation of synthetic ICs needed to be of therapy-grade purity. The assay setup is depicted in Fig. 1D.

**Fig. 1.**
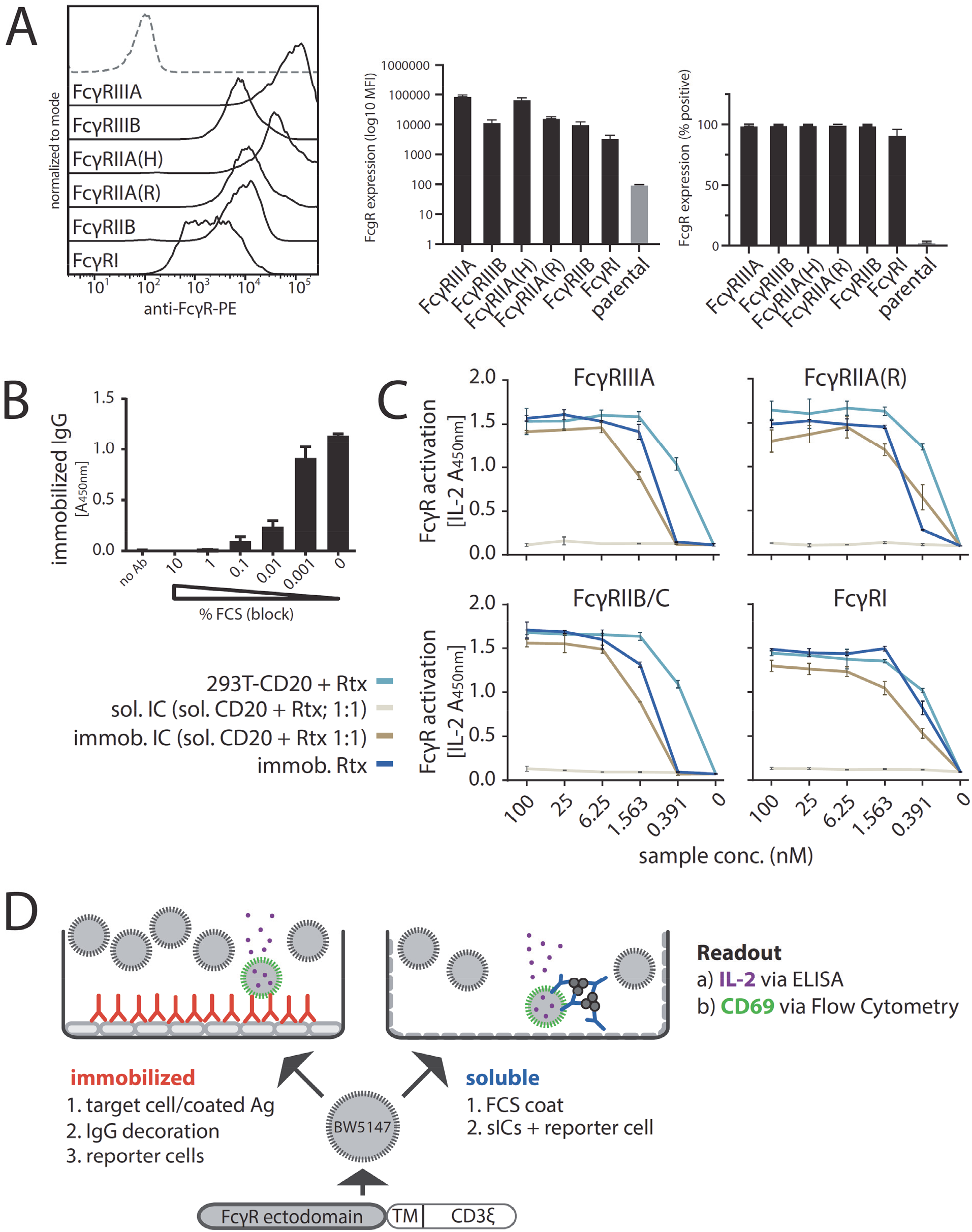
Establishment of a cell-based reporter assay measuring FcγR activation in response to sICs. A) BW5147 reporter cells stably expressing human FcγR-ζ chain chimeras or BW5147 parental cells (grey/dashed) were stained with FcγR specific conjugated mAbs as indicated and measured for surface expression of FcγRs via flow cytometry. B) FCS coating of an ELISA microtiter plate allows for suspension of subsequently added IgG. Plate bound IgG was quantified via ELISA. C) Immobilized IC, immobilized IgG and IgG opsonized cells represent qualitatively similar ligands for FcγRs. Response curves of human FcγRs activated by opsonized cells (293T cells stably expressing CD20 + Rituximab [Rtx]), immobilized IC (rec. soluble CD20 + Rtx) and immobilized IgG (Rtx). sICs formed using monovalent antigen (rec. soluble CD20 + Rtx) do not activate human FcγRs. X-Axis shows sample concentration determined by antibody molarity. Y-Axis shows FcγR activation determined by reporter cell mouse IL-2 production (OD 450nm). Two independent experiments performed in technical duplicates. Error bars = SD. D) Schematic of used assay setups. BW5147 reporter cells expressing chimeric human FcγR receptors express endogenous CD69 or secrete mouse IL-2 in response to FcγR activation by clustered IgG. sICs are generated using mAbs and multivalent antigens. sIC suspension requires pre-blocking of an ELISA plate using PBS supplemented with 10% FCS (FCS coat, grey-dashed).

### Detection of human FcγR activation by multimeric sICs

Next, we generated synthetic sICs from recombinant ultrapure molecules to evaluate the assay. We aimed to avoid the use non-human molecules, misfolded IgG aggregates or IgG-IgG complexes to generate a most native and defined ligand. To date, there are still few commercially available human IgG-antigen pairs that meet both the above mentioned high grade purity requirements while also consisting of at least two antigen monomers. In order to meet these stipulations we focused on three pairs of multivalent antigens and their respective mAbs that were available in required amounts enabling large-scale titration experiments; trimeric rhTNFα:IgG1 infliximab (TNFα:Ifx), dimeric rhVEGFA: IgG1 bevacizumab (VEGFA/Bvz) and dimeric rhIL-5: IgG1 mepolizumab (IL-5/Mpz). As lymphocytes express TNFα-receptors I and II while not expressing receptors for IL-5 or VEGFA, we tested whether the mouse lymphocyte derived BW5147 thymoma reporter cell line is sensitive to high concentrations of rhTNFα. Toxicity testing revealed that even high concentrations of up to 76.75 nM rhTNFα did not affect viability of reporter cells (Fig. S1). Next, we measured the dose-dependent activation of human FcγRs comparing immobilized IgG to sICs (TNFα:Ifx) using the full FcγR reporter cell panel (Fig. 2). Soluble antigen or mAb alone served as negative controls showing no background activation even at high concentrations. Immobilized rituximab (Rtx, human IgG1) and immobilized FcγR-specific mouse mAbs served as positive controls for inter-experimental reference. All FcγRs were activated by immobilized IgG. Only FcγRI failed to respond when reporter cells were incubated with sICs. Both the IL-2 ELISA as well as the CD69 expression read-out gave comparable results. Further, using an IL-2 standard, we were able to quantify FcγR activation (right y-axis). This revealed that the reporter cell lines differ regarding reactivity which did not correlate with receptor expression. Although the low signals for FcγRI might be linked to receptor expression, responsiveness was markedly lower compared to other reporter cell lines (Fig. 1A). Attempts to increase and equalize receptor expression by repeated cell sorting steps failed, indicating that individual receptors only tolerate limited molecule densities on the reporter cell surfaces.

**Fig. 2.**
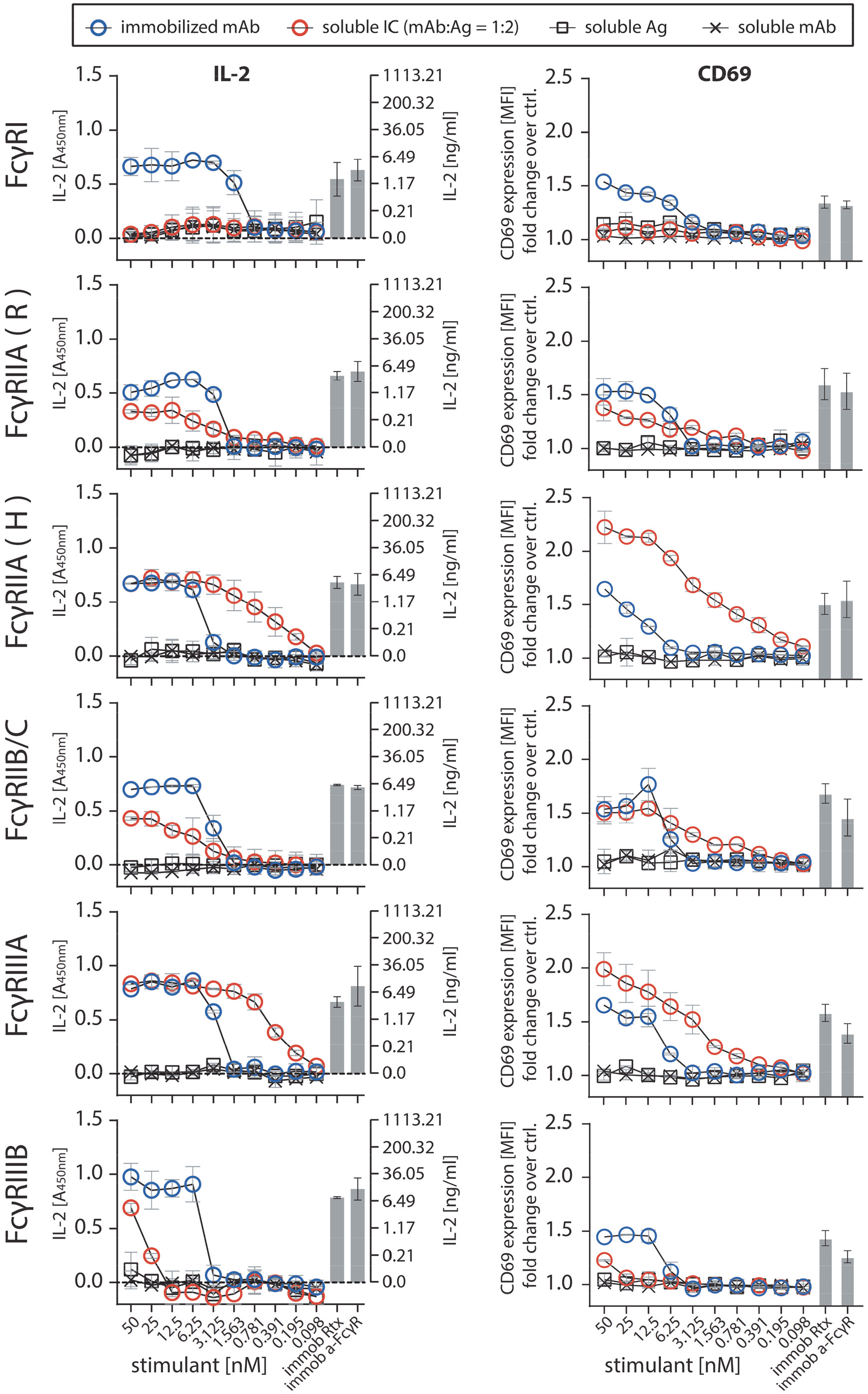
FcγRs are activated by sICs formed from multivalent antigens. Ultra-pure antigen (Ag, TNF-α) mixed with therapy-grade mAb (infliximab, Ifx) was used to generate sICs. X-Axis: concentrations of stimulant expressed as molarity of either mAb or Ag monomer and IC (expressed as mAb molarity) at a mAb:Ag ratio of 1:2. Soluble antigen or soluble antibody alone served as negative controls and were not sufficient to activate human FcγRs. Immobilized IgG (Rtx) or immobilized FcγR-specific mAbs served as positive controls. Two independent experiments performed in technical duplicates. Error bars = SD. Error bars smaller than symbols are not shown. Left panel: IL-2 quantification 16 h after reporter cell activation. Background (blank) was subtracted (dashed line). IL-2 was measured via anti-IL-2 ELISA (A_450nm_) and IL-2 concentrations were calculated from an IL-2 standard. Right panel: Reporter cell CD69 expression 4 h post trigger was measured using flow cytometry. MFI were normalized to untreated cells (ctrl.) and are presented as fold-change increase.

### Evaluation of human FcγR activation by multimeric sICs

The assay proved to be sensitive to sICs in the nanomolar range. Regarding immobilized IgG, the detection limit was between 1 to 3 nM. sICs were detected with the following limits regarding the IL-2 readout: FcγRI – no detection; FcγRIIA(R) – 3 nM; FcγRIIA(H) – 0.2 nM; FcγRIIB/C – 3 nM; FcγRIIIA – 0.2 nM; FcγRIIIB - 25 nM. We observed that sICs and immobilized ICs induce largely different signal strength in individual reporter cell lines. FcγRI and FcγRIIIB were more efficiently activated by immobilized IgG compared to sICs. Conversely, FcγRIIA(H) and FcγRIIIA were more efficiently activated by sICs. FcγRIIB/C showed discrepant results when comparing the IL-2 read-out (16 h) with the CD69 read-out (4 h). Here, it seems that a longer activation leads to a stronger signal on immobilized IgG, while shorter activation slightly favours sIC reactivity. FcγRII(R) looked similar to FcγRIIB/C with a slightly higher response to sICs at low stimulant concentrations. Nevertheless, the response to immobilized IgG was higher for both read-outs at higher concentrations. FcγRIIA(H) and FcγRIIIA proved to be the most sensitive towards sICs stimulation. Notably, the reported superior interactivity of sICs with FcγRII (H) over FcγRII (R) (Shashidharamurthy et al, 2009) was not only confirmed using our assay but we also show that this difference is limited to sIC reactivity and is not seen in immobilized IC reactivity (Fig. 2). Additionally, we measured the response of select reporter cell lines towards sICs of different composition (VEGFA/Bvz and IL-5/Mpz). As these sICs incorporate dimeric antigens, we tested if reporter responses were still comparable (Fig. S2). We observed that responses to sICs were generally lower for FcγRIIA(R) but comparable for FcγRI, FcγRIIB/C and FcγRIIIA. Of note, FcγRI showed slight reactivity towards VEGFA/Bvz sICs. Based on the universal transmembrane and cytosolic part of the FcγR chimeras in our assay we concluded that FcγR ectodomains are intrinsically able to differentiate between different conformations of sICs and immobilized monomeric IgG ligands. To validate the data generated by the reporter assay, we determined FcγRIIIA activation using primary human NK cells isolated from PBMCs of three healthy donors. We chose NK cells as they mostly express only one type of FcγR similar to the reporter system and used IL-5/Mpz sICs as NK cells do not respond to IL-5. Measuring a panel of activation markers and cytokine responses by flow cytometry, we observed a differential activation pattern depending on ICs being soluble or immobilized at equal molarity (Fig. 3A). While MIP1-β responses were comparable between the two triggers, degranulation (CD107a) and TNFα responses showed a trend towards lower activation by sICs compared to immobilized IgG (Mpz). Strikingly, IFNγ responses were significantly weaker when NK cells were incubated with sICs compared to immobilized IgG. Next, in order to confirm this to be due to specific activation of FcγRIIIA, we changed the sIC setup by generating reverse-orientation sICs consisting of human FcγR-specific mouse mAbs and goat-anti-mouse IgG F(ab)_2_ fragments (Fig. S3A). NK cell activation by reverse sICs was compared to NK cell activation by immobilized FcγR specific mAbs. This confirmed our previous observations. As in roughly 10% of the population NK cells express FcγRIIC (Anania et al, 2019; Breunis et al, 2008; Lisi et al, 2011; Metes et al, 1998), we also tested reverse sIC activation using an FcγRII specific mAb. As we did not observe an FcγRII-mediated response, we conclude that FcγRIIC expression did not play a role in our experiments (Fig. S3B). Importantly, these experiments validate that all our experimentally synthesized sICs readily activate primary NK cells and induce immunological effector functions.

**Fig. 3.**
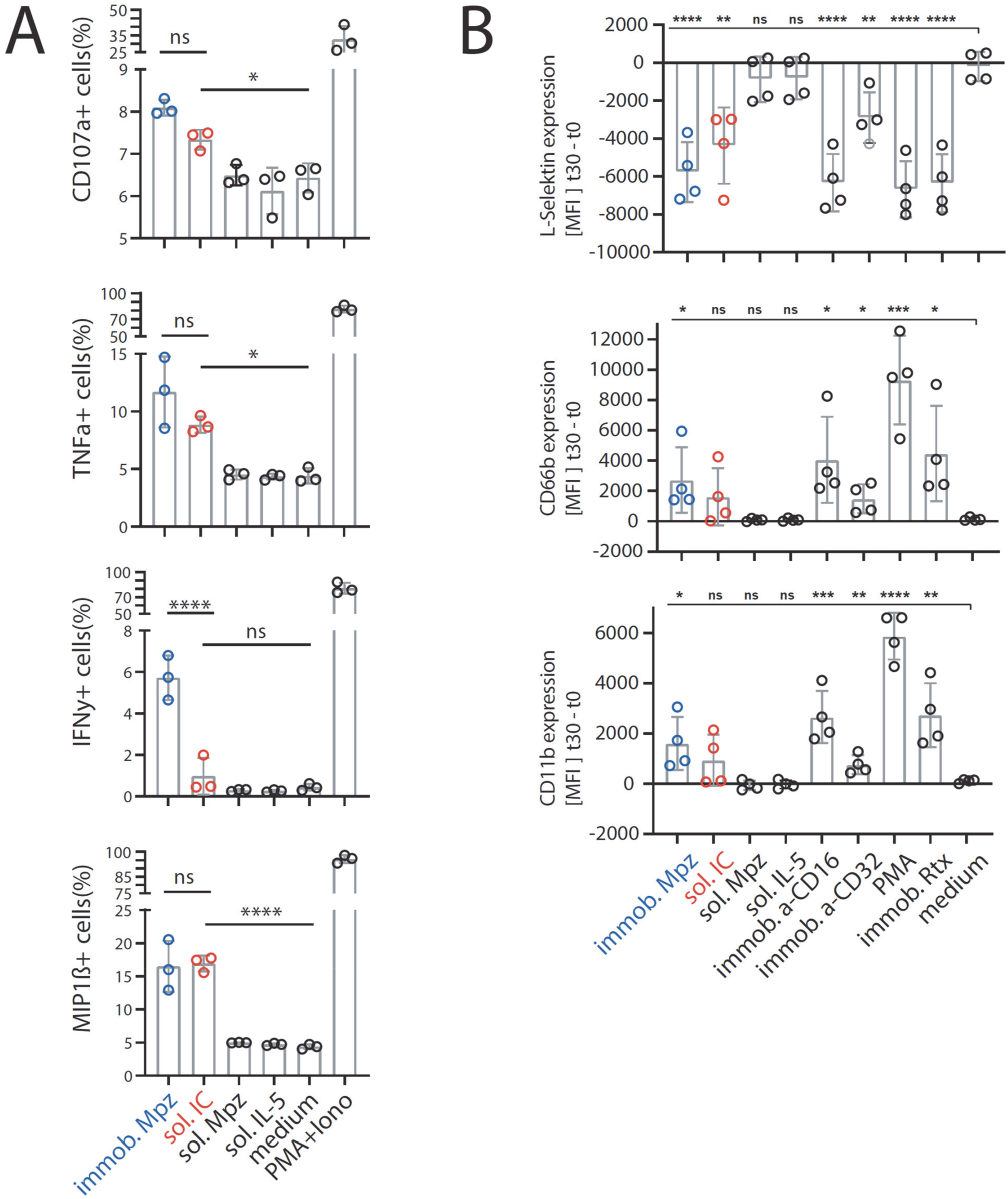
The FcγR-dependent activation pattern of primary NK cells or primary neutrophils depends on IC solubility. A) Negatively selected primary NK cells purified from PBMCs of three healthy donors were tested for activation markers using flow cytometry. NK cells were incubated with immobilized IgG (mepolizumab, Mpz), soluble IC (Mpz:IL-5 = 1:1), soluble Mpz or soluble IL-5 (all at 200 nM, 10^6^ cells). Incubation with PMA and Ionomycin (Iono) served as a positive control. Incubation with medium alone served as a negative control. Means of technical duplicates. Error bars = SD. One-way ANOVA (Tukey); *p<0.05, **p<0.01, ***p<0.001, ****p<0.0001. B) Negatively selected primary neutrophils purified from whole blood of four healthy donors were tested for adhesion and activation markers using flow cytometry. Neutrophils were incubated with immobilized IgG (Mpz), soluble IC (Mpz:IL-5 = 1:1), soluble Mpz or soluble IL-5 (all at 200 nM, 2*10^5^ cells). Incubation with PMA or immobilized rituximab served as positive controls. Incubation with medium served as a negative control. Immobilized FcγRII and FcγRIII specific mAbs served as functional controls. Mean florescence intensity (MFI) values at t=30 minutes of incubation are presented as increase over t=0 min. Means of technical duplicates. Error bars = SD. Two-way ANOVA compared to medium (Dunnett); *p<0.05, **p<0.01, ***p<0.001, ****p<0.0001.

Next, primary neutrophils isolated from whole blood samples of four individual donors were analysed, measuring upregulation of CD11B, CD66B and shedding of L-selectin as markers of immune complex mediated adhesion and activation (Ilton et al, 1999; Khawaja et al, 2019; Lard et al, 1999; Zarbock & Ley, 2009) (Fig. 3B). Again, immobilized IgG and sICs activated primary neutrophils with different efficiency. While both, sICs and immobilized IgG, strongly induced the shedding of L-selectin, the upregulation of CD11B and CD66B showed a tendency towards lower activation by sICs compared to immobilized IgG. As neutrophils express both FcγRIIA and FcγRIIIB, we also individually activated these receptors on neutrophils using immobilized FcγR-specific mAbs. This revealed that neutrophil activation by FcγRs is mostly driven by FcγRIIIB. In light of our previous tests (Fig. 2), this explains the reduced activation by sICs. Taken together, we conclude that primary cells differentiate between opsonized targets and sICs via inherent features of individual FcγRs as well as by the co-expression of FcγRs with different sensitivity towards sICs.

### Measurement of FcγR activation in response to the molecular size of sICs

We observed that the dimeric CD20:Rtx molecule complex completely failed to trigger FcγR activation (Fig. 1C) while potentially larger sICs, based on multimeric antigens, showed an efficient dose-dependent FcγR activation (Fig. 2, S2). In order to determine whether FcγR signalling responds to changes in sIC size, we cross-titrated amounts of antibody (mAb, infliximab, Ifx) and antigen (Ag, rhTNFα). Specifically, the reporter cells were incubated with sICs of varying mAb:Ag ratio by fixing one parameter and titrating the other. According to the Heidelberger-Kendall precipitation curve (Heidelberger & Kendall, 1929), sIC size depends on the mAb:Ag ratio. sICs of varying sizes result from an excess of either antigen or antibody, leading to the formation of smaller complexes compared to the large molecular complexes formed at around equal molarity. Presumed changes in sIC size were quantified using asymmetrical flow-field flow fractionation (AF4) (Fig. 4A and Table S1). Fig. S4 shows a complete run of an exemplary analysis. Af4 analysis revealed a sIC mean molecular weight of approximately 2130 kDa at a 1:3 ratio (Ifx/TNF-α) with sICs getting smaller with increasing excess of either antigen or antibody, recapitulating a Heidelberger-Kendall-like curve. Incubation of the FcγR reporter cells with sICs of varying size indeed shows that the assay is highly sensitive to changes in sIC size (Fig. 4B). Accordingly, FcγRs showed the strongest responses at mAb:Ag ratios of approximately 1:3. Next, we validated the accuracy of our reporter cell data by subjecting primary human NK cells to the same variation of mAb:Ag stoichiometry. NK cells from three individual donors were measured for MIP1-β upregulation in response to synthetic sICs of varying size and composition (Fig. 4C). Indeed, primary NK cells equally responded to sIC size at the same nanomolar range of stimulating ligand, confirming that the reporter system accurately measures immune cell responses to sICs. Convincingly, NK cell responses to sICs generated from trimeric antigen (TNFα) peaked at a different mAb:Ag ratio compared to NK cell responses to sICs generated from dimeric antigens (IL-5 and VEGFA). TNFα and VEGFA contribute to the activation of resting NK cells, thus leading to higher MIP1-β positivity when NK cells are incubated in the presence of excess antigen. As NK cells do not express IL-5 receptor, this effect is not observed in the presence of excess IL-5. The data reveal a direct correlation between sIC dimension and effector responses. Conversely, when changing antibody concentrations using fixed amounts of antigen, a consistent reduction of NK cell activation is observed in the presence of excess IgG for all three mAb:Ag pairs.

**Fig. 4.**
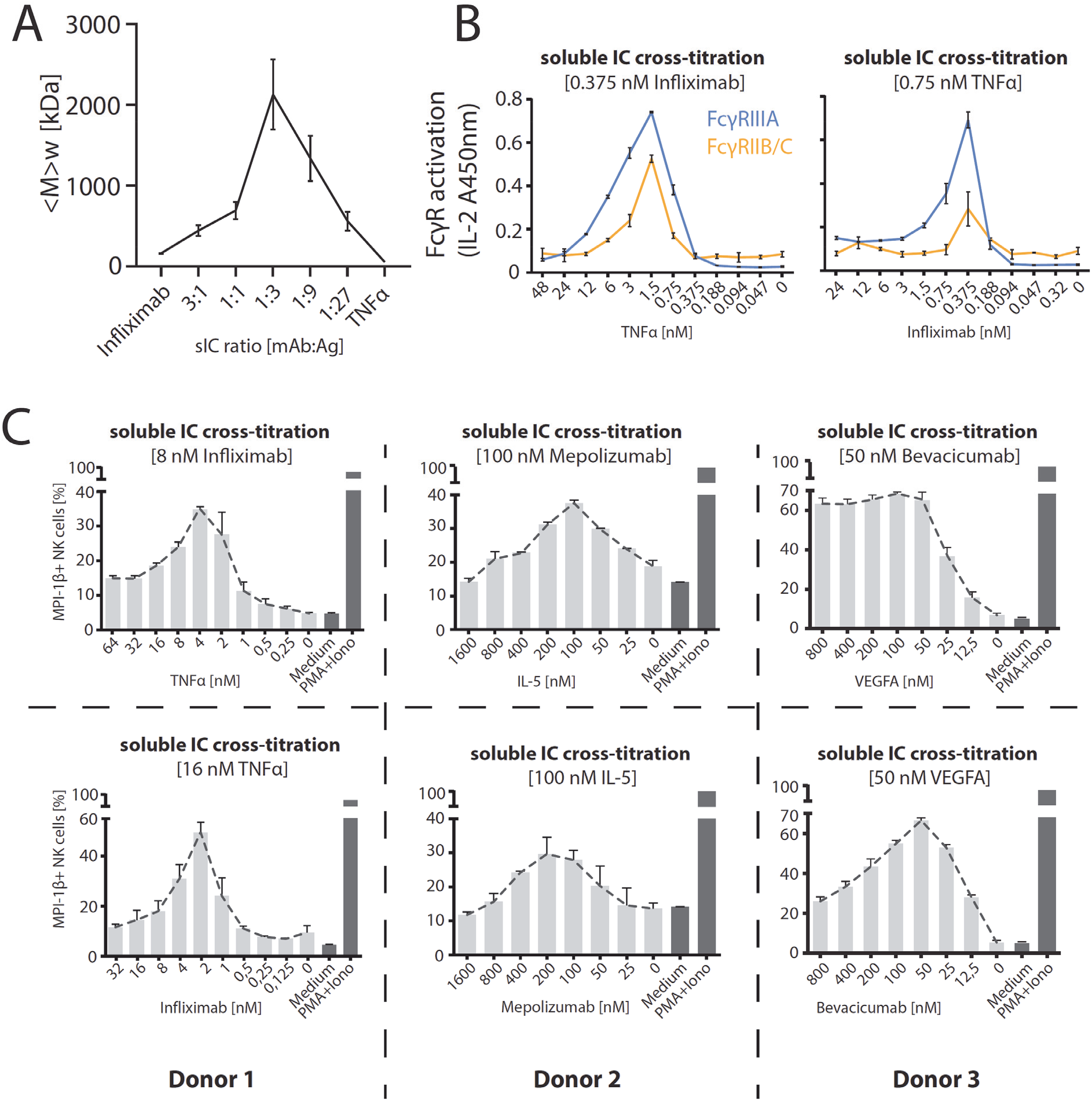
FcγRIIB/C and FcγR-IIIA respond to sIC size reproducing a Heidelberger-Kendall like precipitation curve. A) infliximab (mAb) and rhTNFα (Ag) were mixed at different ratios (17 µg total protein, calculated from monomer molarity) and analysed via AF4. sIC size is maximal at a 1:3 ratio of mAb:Ag and reduced when either mAb or Ag are given in excess. <M>_w_ = mass-weighted mean of the molar mass distribution. Three independent experiments. Error bars = SD. Data taken from Table S1. One complete run analysis is shown in Fig. S2. B) sICs of different size were generated by cross-titration according to the AF4 determination. Reporter cells were incubated with fixed amounts of either mAb (infliximab, left) or Ag (rhTNFα, right) and titrated amounts of antigen or antibody, respectively. X-Axis shows titration of either antigen or antibody, respectively (TNFα calculated as monomer). Two independent experiments performed in technical duplicates. Error bars = SD. C) Purified primary NK cells from three different donors were incubated with cross-titrated sICs as in A. NK cells were measured for MIP-1β expression (% positivity). Incubation with PMA and Ionomycin served as a positive control. Incubation with medium alone served as a negative control. Measured in technical duplicates. Error bars = SD.

### Quantification of sIC bioactivity in sera of SLE patients

In order to apply the assay to a clinically relevant setting associated with the occurrence of sICs, we measured circulating sICs present in the serum of SLE patients with varying disease activity. Sera from 4 healthy donors and 25 SLE patients were investigated for FcγRIIIA and FcγRIIB/C activation to compare an activating and an inhibitory receptor. Reporter cells readily secreted mIL-2 in response to patient sera in a dose-dependent manner (Fig. 5A), which was not the case when sera from healthy controls were tested. We confirm that FcγRIIIA and FcγRIIB/C activation depends on the presence of serum sICs by comparing the bioactivity of patient serum before and after polyethylene glycol (PEG) precipitation which is known to deplete sICs (Lux et al, 2013) (Fig. 5B). Next, we calculated the area under the curve (AUC) values for all 25 SLE patient titrations and normalized them to the AUC values measured for healthy individuals. The resulting index values were then correlated with established biomarkers of SLE disease activity, being anti-dsDNA titers (α-dsDNA) and concentrations of the complement cleavage product C3d (Fig. 5C). We observed a significant correlation between our FcγRIIIA activation index values and both disease activity markers (p=0.0465 and p=0.0052, respectively). FcγRIIB/C activation showed no significant correlation with either biomarker. We assume these interrelations may be due to the influence of IgG sialylation found to be reduced in active SLE (Vuckovic et al, 2015). Generally, de-sialylation of IgG leads to stronger binding by the activating receptors FcγRI, FcγRIIA and FcγRIII while it reduces the binding affinity of the inhibitory FcγRIIB (Kaneko et al, 2006). In practice, our assay allows the detection and quantification of clinically relevant sICs in sera from SLE patients as shown here or in synovial fluid of rheumatoid arthritis patients (Zhao et al, 2021).

**Fig. 5.**
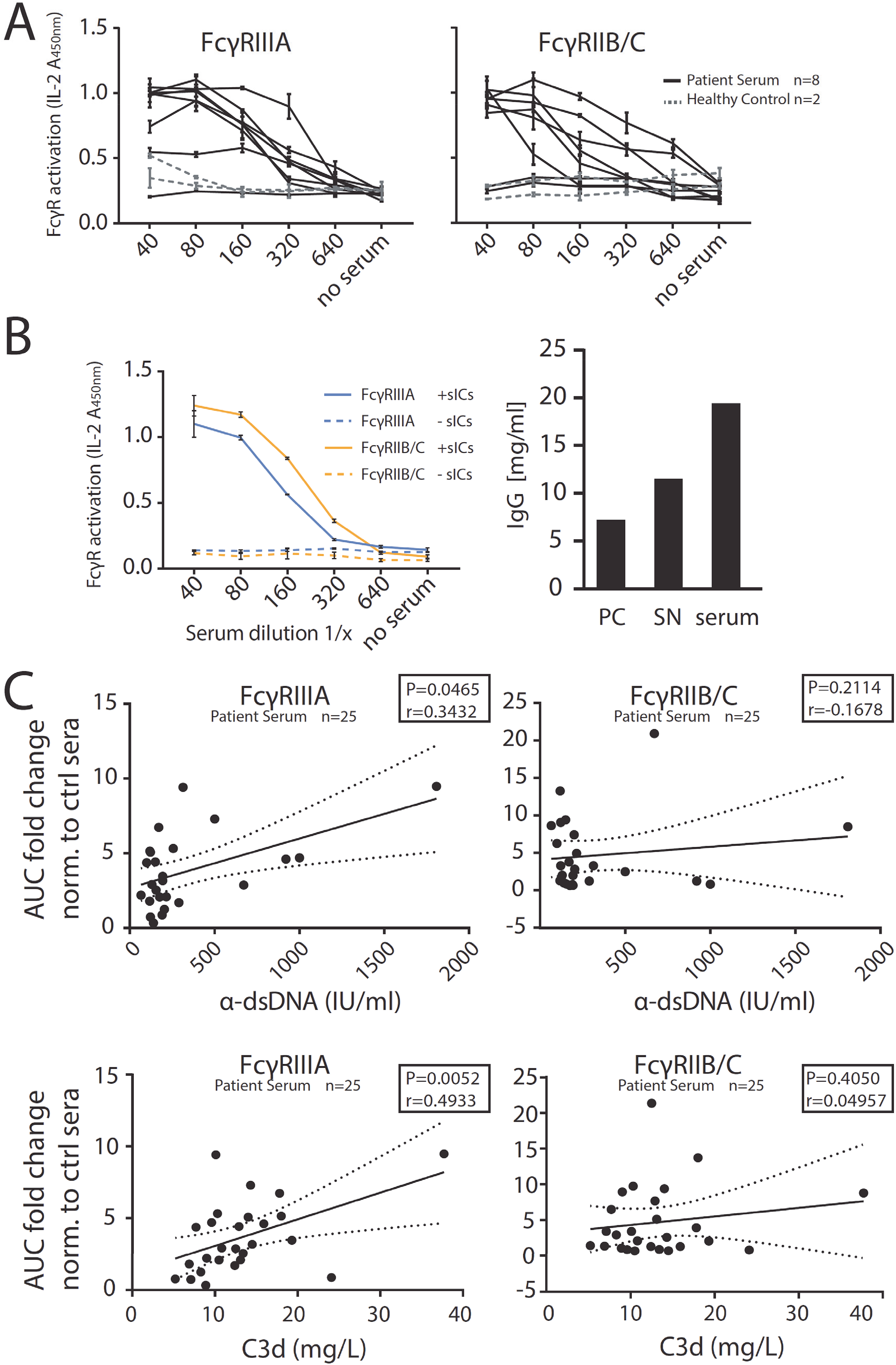
The reporter assay enables quantification of serum-derived sICs from SLE patients. Serum derived sIC from systemic lupus erythematosus (SLE) patients activate human FcγR reporter cells. 25 patients and 4 healthy control individuals were separated into three groups for measurement. A) Experiments shown for an exemplary group of 8 SLE patients and two healthy individuals. Dose-dependent reactivity of FcγRs IIIA and IIB/C was observed only for SLE patient sera and not for sera from healthy individuals. One exemplary experiment performed in technical duplicates. Error bars = SD. B) Activation of FcγRs IIB/C and IIIA by patient serum is mediated by serum derived sICs. Patient serum samples were depleted of sICs by PEG precipitation and the supernatant (SN) was compared to untreated serum regarding FcγR activation (left). One experiment performed in technical duplicates. Error bars =SD. IgG concentration in the precipitate (PC), supernatant (SN) or unfractionated serum respectively is shown in the bar graph (right). C) FcγR activation data from A was compared to conventional SLE disease markers (α-dsDNA levels indicated as IU/ml or C3d concentrations indicated as mg/L). FcγR activation from a dose-response curve as in A was calculated as area under curve (AUC) for each SLE patient (n=25) or healthy individual (n=4) and expressed as fold change compared to the healthy control mean. SLE patients with α-dsDNA levels below 50 IU/ml and C3d values below 6 mg/L were excluded. One-tailed Spearman’s.

### Assay application to in vivo mouse models of lupus and arthritis

BW5147 reporter cells stably expressing chimeric mouse as well as rhesus macaque FcγRs have already been generated using the here described methodology (Kolb et al, 2019; Van den Hoecke et al, 2017). Next we aimed to translate the assay to clinically relevant mouse models. FcγR reporter cells expressing chimeric mouse FcγRs were incubated with sera from lupus (NZB/WF1)(Dubois et al, 1966) or arthritis (K/BxN)(Kouskoff et al, 1996) mice with symptomatic disease. We chose to determine the stimulation of the activating receptors, mFcγRIII and mFcγRIV. Incubation with synthetic sICs generated from rhTNFα and mouse-anti-hTNFα IgG1 showed both of the reporters to be equally responsive to sICs (Fig. 6A). Parental BW5147 cells expressing no FcγRs served as a control. The sera of three mice per group were analysed and compared to sera from wildtype C57BL/6 mice, which served as a healthy control. C57BL/6 mice were chosen, as K/BxN or NZB/WF1 mice show temporal variability in disease onset and presymptomatic phase. We consistently detected mFcγR activation by sera from K/BxN or NZB/WF1 but not healthy C57BL/6 mice (Fig. 6B). While the mFcγRIII responses were generally high and similar between K/BxN and NZB/WF1 mice, mFcγRIV responsiveness tended to be lower and individually more variable. Altogether, the assay enables the reliable detection of sICs in sera of mice with immune-complex mediated diseases making it a promising novel research tool to study the role of sIC formation and FcγR activation in preclinical mouse models.

**Fig. 6.**
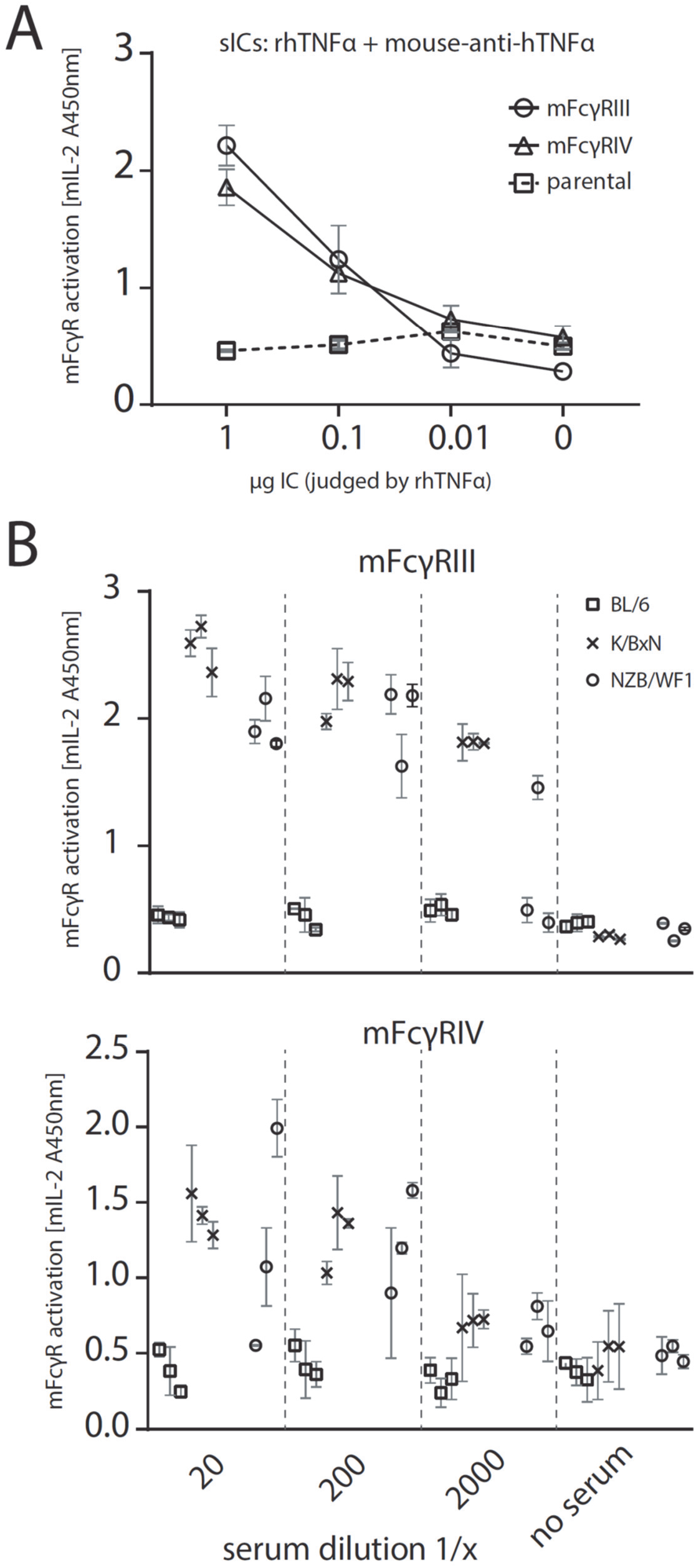
The reporter assay can be applied to mouse models of autoimmune disease. A) Reporter cells expressing mFcγRIII, mFcγRIV or parental BW5147 cells were incubated with titrated amounts of synthetic sICs generated from rhTNFα and mouse-anti-hTNFα at a 1:1 ratio by mass. One experiment performed in technical duplicates. Error bars = SD. B) Titrations of 3 mouse sera per group (C57BL/6, K/BxN or NZB/WF1) were incubated with mFcγR reporter cells and FcγR activation was assessed as described above. Sera from BL/6 mice served as negative control. Two independent experiments in technical duplicates. Error bars = SD.

## Discussion

In this study we established, validated and applied a new assay system that is able to selectively detect soluble multimeric immune complexes as discrete ligands of FcγRs. Our system is sensitive to the size, concentration and composition of sICs. The assay is scalable and supports measurement with human and mouse FcγRs. It provides two readouts suitable for high-throughput analysis: fast CD69 surface expression and quantifiable IL-2 secretion.

### A novel assay for the quantification of individual FcγR activation by experimental and clinical sICs

Our methodology provides a comprehensive system, supporting the assessment of essentially all FcγRs, which presents an advantage over previously developed sIC detection and FcγR activation assays (Aoyama et al, 2019; Cheng et al, 2014; Hsieh et al, 2017; Stopforth et al, 2018; Szittner et al, 2016; Tada et al, 2014). In contrast to currently available commercial assays detecting sICs by C1q-CIC or C3d ELISA in the micromolar range, our assay measures overall sIC bioactivity in the nanomolar range and has a sole specificity for IgG sICs. The new approach presents with hands on technical advances as it allows for the measurement of small or large amounts of samples by a relatively simple *in vitro* assay with high-throughput potential. Favourably, BW5147 reporter cells are largely inert to human cytokines, which provides a key advantage to measure their responsiveness after contact with human samples. Our pilot study demonstrates that sIC-mediated FcγRIIIA activation correlates with conventional SLE disease markers. This is of great value as a recent analysis shows that circulating sICs and IL-6 can predict SLE activity with the higher accuracy compared to conventional clinical SLE biomarkers (Thanadetsuntorn et al, 2018). However, circulating immune complexes in this study were determined using a commercial C1q-binding ELISA, lacking information on immune cell bioactivity of the measured sICs. Our assay should therefore be explored as an addition to the clinician’s toolbox which may allow better disease management. Due to the scalability and high-throughput readouts, the assay can also be of use for larger prospective clinical studies in patients with autoimmune diseases such as SLE or rheumatoid arthritis, where circulating sICs have long been shown to crucially contribute to tissue damage and disease manifestations (Koffler et al, 1971; Levinsky, 1978; Levinsky et al, 1977; Nydegger & Davis, 1980; Zubler et al, 1976). Disease-associated, endogenous sICs can also be formed from multimeric viral and bacterial structural proteins generated during infection (Briant et al, 1996; Oh et al, 1992; Vuitton et al, 2020), where circulating sICs strongly impact pathogenesis (Madalinski et al, 1991; Wang & Ravetch, 2015).

### Dynamic sIC size measurement and monitoring of bioactivity in sIC-associated diseases

The new sIC approach allowed for a simultaneous functional and biophysical assessment of the paradigmatic Heidelberger-Kendall precipitation curve (Heidelberger & Kendall, 1929; Heidelberger & Kendall, 1935). While previous work already revealed that large and small sICs differentially impact IL-6 production in PBMCs (Lux et al, 2013), the dynamics of FcγR activation resulting from constant changes in sIC size have not been explored systematically and lacked resolution of defined FcγR types. We analyzed synthetic sICs formed by highly pure recombinant components via AF4. Our data document that sIC size is indeed governed by antibody:antigen ratios covering a wide range of sizes up to several megadaltons. In the presence of increasing amounts of antibody or antigen deviating from an optimal antibody:ratio, sIC size steadily decreases. Further, by the measurement of FcγR activation we now translate physical sIC size directly to a simple but precise biological read-out. In doing so, we show that sIC size essentially tunes FcγR activation on and off. Thus, our new test system can not only contribute to the functional detection and quantification of clinically relevant sICs but also provides a starting point on how to avoid pathological consequences by influencing the sIC size, for example by administering and monitoring of therapeutic antibodies or recombinant antigens in controlled amounts, thus becoming relevant in clinical pharmacokinetics.

### Limitations of the reporter system and conclusions

There is a wide range of factors, regulating and influencing the sIC-FcγR interaction. These include Fcγ-FcγR binding affinity and avidity (Koenderman, 2019), IgG subclass, IgG glycan profiles and genetic polymorphism (Bruhns et al, 2009; Pincetic et al, 2014; Plomp et al, 2017; Vidarsson et al, 2014), stoichiometry of antigen-antibody-ratio (Berger et al, 1996; Lux et al, 2013; Pierson et al, 2007), FcγR clustering patterns (Patel et al, 2019), downstream signaling (Bournazos et al, 2017; Getahun & Cambier, 2015) and the interaction of FcγR with other receptors (Douek et al, 2009; Ortiz-Stern & Rosales, 2003; Urbaczek et al, 2014; van Egmond et al, 2015; Vanderbruggen et al, 1994). Our assay is sensitive to amount, size and glycosylation of sICs and can readily be adapted to include more FcγR genotypes and polymorphisms by generation of additional reporter cell lines.

The major advancements of this reporter system include i) a high accuracy and resolution regarding FcγR type-specific activation compared to traditional indirect assessment via affinity measurements, ii) a scalable and quantifiable assay providing flexible high-throughput readouts in the nanomolar range, iii) an sIC size sensitive reporter system and iv) a comprehensive panel including all human FcγRs. In practice, the platform is suitable to be implemented into small- or large-scale screening setups in research as well as routine laboratories. Prospectively, the reporter cell approach allows for future adaptation as the cells can be equipped with alternative reporter modules to optimize the methodology for specific applications.

## Materials and Methods

### Cell culture

All cells were cultured in a 5% CO_2_ atmosphere at 37°C. BW5147 mouse thymoma cells (BW, kindly provided by Ofer Mandelboim, Hadassah Hospital, Jerusalem, Israel) were maintained at 3×10^5^ to 9×10^5^ cells/ml in Roswell Park Memorial Institute medium (RPMI GlutaMAX, Gibco) supplemented with 10% (vol/vol) fetal calf serum (FCS, Biochrom), sodium pyruvate (1x, Gibco) and β-mercaptoethanol (0.1 mM, Gibco). 293T-CD20 (kindly provided by Irvin Chen, UCLA (Morizono et al, 2010)) were maintained in Dulbecco’s modified Eagle’s medium (DMEM, Gibco) supplemented with 10% (vol/vol) FCS.

### BW5147 cell flow cytometry

BW5147 cells were harvested by centrifugation at 900 g and RT from the suspension culture. 1×10^6^ cells were stained with PE-conjugated anti-human FcγR mAbs (BD) or a PE-TexasRed-conjugated human IgG-Fc fragment (Rockland) for 1h at 4°C in PBS/3%FCS. After 3 washing steps in PBS/3%FCS, the cells were transferred to Flow cytometry tubes (BD) and analysed using BD LSR Fortessa and FlowJo (V10) software. Cells sorting was performed at the Lighthouse core facility of the University Hospital Freiburg using receptor staining (BD Pharmingen, PE-conjugated).

### Lentiviral transduction

Lentiviral transduction of BW5147 cells was performed as described previously (Halenius et al, 2011; Kolb et al, 2019; Van den Hoecke et al, 2017). In brief, chimeric FcγR-CD3ζ constructs (Corrales-Aguilar et al, 2013) were cloned into a pUC2CL6IPwo plasmid backbone. For every construct, one 10-cm dish of packaging cell line at roughly 70% density was transfected with the target construct and two supplementing vectors providing the VSV gag/pol and VSV-G-env proteins (6 µg of DNA each) using polyethylenimine (22.5 µg/ml, Sigma) and Polybrene (4 µg/ml; Merck Millipore) in a total volume of 7 ml (2 ml of a 15-min-preincubated transfection mix in serum-free DMEM added to 5 ml of fresh full DMEM). After a medium change, virus supernatant harvested from the packaging cell line 2 days after transfection was then incubated with target BW cells overnight (3.5 ml of supernatant on 10^6^ target cells), followed by expansion and pool selection using complete medium supplemented with 2 µg/ml of puromycin (Sigma) over a one week culture period.

### human IgG suspension ELISA

1 µg of IgG1 (rituximab in PBS, 50 µl/well) per well was incubated on a 96well microtiter plate (NUNC Maxisorp) pre-treated (2h at RT) with PBS supplemented with varying percentages (v/v) of FCS (PAN Biotech). IgG1 bound to the plates was detected using an HRP-conjugated mouse-anti-human IgG mAb (Jackson ImmunoResearch).

### Recombinant antigens and monoclonal antibodies to form sICs

Recombinant human (rh) cytokines TNF, IL-5, and VEGFA were obtained from Stem Cell technologies. Recombinant CD20 was obtained as a peptide (aa141-188, Acc# P11836) containing the binding region of rituximab (Creative Biolabs). FcγR-specific mAbs were obtained from Stem Cell technologies (CD16: clone 3G8; CD32: IV.3). Reverse sICs were generated from these receptor-specific antibodies using goat-anti-mouse IgG F(ab)_2_ fragments (Invitrogen) in a 1:1 ratio. Pharmaceutically produced humanized monoclonal IgG1 antibodies infliximab (Ifx), bevacizumab (Bvz), mepolizumab (Mpz) and rituximab (Rtx) were obtained from the University Hospital Pharmacy Freiburg. Mouse anti-hTNFα (IgG2b, R&D Systems, 983003) was used to generate sICs reactive with mouse FcγRs. sICs were generated by incubation of antigens and antibodies in reporter cell medium or PBS for 2 h at 37°C.

### FcγR receptor activation assay

FcγR activation was measured adapting a previously described cell-based assay (Corrales-Aguilar et al, 2014; Corrales-Aguilar et al, 2013). The assay was modified to measure FcγR activation in solution. Briefly, 2×10^5^ mouse BW-FcγR (BW5147) reporter cells were incubated with synthetic sICs or diluted serum in a total volume of 100 µl for 16 h at 37°C and 5% CO_2_. Incubation was performed in a 96-well ELISA plate (Nunc maxisorp) pre-treated with PBS/10% FCS (v/v) for 1 h at 4°C. Immobilized IgG was incubated in PBS on the plates prior to PBS/10% FCS treatment. After 4h incubation, surface mouse CD69 expression was measured using a high throughout sampler (HTS)-FACS. Reporter cell mouse IL-2 secretion was quantified after 16 h of incubation via anti-IL-2 ELISA as described earlier (Corrales-Aguilar et al, 2013).

### High throughout sampler flow cytometry (HTS-FACS)

After 4h of stimulation, 1×10^5^ BW5147 reporter cells were stained with APC-conjugated anti-mCD69 (Biolegend; CD69: H1.2F3; 1:100) for 30min at 4°C in PBS/3%FCS. Cells were transferred to a U Form 96well Microplate (Greiner 650101) and analysed by flow cytometry (BD Fortessa). High Throughput mode was designed within BD FACSDiva software using HTS mode with the following parameters: sample flow rate 2µl/s, sample volume 10µl, mixing volume 50µl, mixing speed 200µl/s, number of mixes 2 cycles and wash volume 200µl.

### BW5147 toxicity test

Cell counting was performed using a Countess II (Life Technologies) according to supplier instructions. Cell toxicity was measured as a ratio between live and dead cells judged by trypan blue staining over a 16 h time frame in a 96well format (100 µl volume per well). BW5147 cells were mixed 1:1 with trypan blue (Invitrogen) and analysed using a Countess II. rhTNFα was diluted in complete medium.

### NK cell activation flow cytometry

PBMC were purified from donor blood using Lymphocyte separation Media (Anprotec). Blood draw and PBMC purification from donors was approved by vote 474/18 (ethical review committee, University of Freiburg). Primary NK cells were separated from donor PBMCs via magnetic bead negative selection (Stem Cell technologies) and NK cell purity was confirmed via staining of CD3 (Biolegend, clone HIT3a), CD16 (Biolegend, clone 3G8) and CD56 (Miltenyi Biotec, clone AF12-7H3). 96well ELISA plates (Nunc Maxisorp) were pre-treated with PBS/10% FCS (v/v) for 1 h at 4°C. NK cells were stimulated in pre-treated plates and incubated at 37°C and 5% CO_2_ for 4 h. Golgi Plug and Golgi Stop solutions (BD) were added as suggested by supplier. CD107a (APC, BD, H4A3) specific conjugated mAb was added at the beginning of the incubation period. Following the stimulation period, MIP-1β (PE, BD Pharmingen), IFNγ (BV-510, Biolegends, 4SB3) and TNFα (PE/Cy7, Biolegends, MAB11) production was measured via intracellular staining Cytokines (BD, CytoFix/CytoPerm, Kit as suggested by the supplier). 50 ng/ml PMA (InvivoGen) + 0.5 µM Ionomycin (InvivoGen) were used as a positive stimulation control for NK cell activation. After 3 washing steps in PBS/3%FCS, the cells were transferred to Flow cytometry tubes (BD) and analysed using a BD FACS Fortessa and FlowJo (V10) software.

### Neutrophil adhesion and activation flow cytometry

Human primary neutrophil granulocytes were isolated from whole blood of healthy donors via magnetic bead negative selection (Stemcell #19666). 96well ELISA plates (Nunc Maxisorp) were pre-treated with PBS/10% FCS (v/v) for 1 h at 4°C. Per reaction, 2×10^5^ cells/ml neutrophils were stimulated with ICs in Roswell Park Memorial Institute medium (RPMI GlutaMAX, Gibco) supplemented with 10% (vol/vol) fetal calf serum (FCS, Biochrom) and incubated at 37°C and 5% CO_2_ for 30 min. Adhesion and activation markers of neutrophils were measured by surface staining of CD11B (APC, Biolegend, ICRF44), CD66B (FITC, Stemcell, G10F5) and L-selectin (PE, Biolegend, DREG-56)(Ilton et al, 1999; Khawaja et al, 2019; Lard et al, 1999; Veen et al, 1998). Cells were then analysed by flow cytometry. FcγRII or FcγRIII cross-linking controls were performed by immobilization of receptor specific mAbs (Stem cell technologies, IV.3 and 3G8) before the ELISA plate was blocked.

### Asymmetric flow field flow fractionation (AF4)

The AF4 system consisted of a flow controller (Eclipse AF4, Wyatt), a MALS detector (DAWN Heleos II, Wyatt), a UV detector (1260 Infinity G1314F, Agilent) and the separation channel (SC channel, PES membrane, cut-off 10 kDa, 490 µm spacer, wide type, Wyatt). Elution buffer: 1.15 g/L Na_2_HPO_4_ (Merck), 0.20 g/L NaH_2_PO_4_ x H_2_O (Merck), 8.00 g/L NaCl (Sigma) and 0,20 g/L NaN_3_ (Sigma), adjusted to pH 7.4, filtered through 0.1 µm. AF4 sequence (Vx = cross flow in mL/min): (a) elution (2 min, Vx: 1.0); (b) focus (1 min, Vx: 1.0), focus + inject (1 min, Vx: 1.0, inject flow: 0.2 mL/min), repeated three times; (c) elution (30 min, linear Vx gradient: 1.0 to 0.0); (d) elution (15 min, Vx: 0.0); (e) elution + inject (5 min, Vx: 0.0). A total protein mass of 17±0.3 µg (Ifx, rhTNFα or ICs, respectively) was injected. The eluted sample concentration was calculated from the UV signal at 280 nm using extinction coefficients of 1.240 mL/(mg cm) or 1.450 mL/(mg cm) in the case of TNFα or Ifx, respectively. For the ICs, extinction coefficients were not available and difficult to calculate as the exact stoichiometry is not known. An extinction coefficient of 1.450 mL/(mg cm) was used for calculating the molar masses of all ICs. Especially in the case of ICs rich in TNFα, the true coefficients should be lower, and the molar masses of these complexes are overestimated by not more than 14 %. The determined molar masses for TNFα-rich complexes are therefore biased but the observed variations in molar mass for the different ICs remain valid. The mass-weighted mean of the distribution of molar masses for each sample was calculated using the ASTRA 7 software package (Wyatt).

### SLE patient cohort

Sera from patients with SLE were obtained from the Immunologic, Rheumatologic Biobank (IR-B) of the Department of Rheumatology and Clinical Immunology. Biobanking and the project were approved by the local ethical committee of the University of Freiburg (votes 507/16 and 624/14). All patients who provided blood to the biobank had provided written informed consent. Ethical Statement: The study was designed in accordance with the guidelines of the Declaration of Helsinki (revised 2013). Patients with SLE (*n* = 25) and healthy controls (*n* = 4) were examined. All patients met the revised ACR classification criteria for SLE. Disease activity was assessed using the SLEDAI-2K score. C3d levels were analyzed in EDTA plasma using rocket double decker immune-electrophoresis with antisera against C3d (Polyclonal Rabbit Anti-Human C3d Complement, Agilent) and C3c (Polyclonal Rabbit Anti-Human C3c Complement Agilent) as previously described (Rother et al, 1993). Anti-human dsDNA antibodies titers were determined in serum using an anti-dsDNA IgG ELISA kit (diagnostik-a GmbH).

### Patient serum IC precipitation

For polyethylene glycol (PEG) precipitation human sera were mixed with PEG 6000 (Sigma-Aldrich) in PBS at a final concentration of 10% PEG 6000. After overnight incubation at 4°C, ICs were precipitated by centrifugation at 2000 x g for 30 min at 4 °C, pellets were washed once with PEG 6000 and then centrifuged at 2000 x g for 20 min at 4 °C. Supernatants were harvested and precipitates re-suspended in pre-warmed PBS for 1 h at 37 °C. IgG concentrations of serum, precipitates and supernatants obtained after precipitation were quantified by Nanodrop (Thermo Scientific™) measurement.

### Mice and Models

Animal experiments were approved by the local governmental commission for animal protection of Freiburg (Regierungspräsidium Freiburg, approval no. G16/59 and G19/21). Lupus-prone (NZBxNZW)F1 mice (NZB/WF1) were generated by crossing NZB*/*BlNJ mice with NZW/LacJ mice, purchased from The Jackson Laboratory. KRNtg mice were obtained from F. Nimmerjahn (Universität Erlangen-Nürnberg) with the permission of D. Mathis and C. Benoist (Harvard Medical School, Boston, MA), C57BL/6 mice (BL/6) and NOD/ShiLtJArc (NOD/Lt) mice were obtained from the Charles River Laboratories. K/BxN (KRNtgxNOD)F1 mice (K/BxN) were obtained by crossing KRNtg mice and NOD/Lt mice. All mice were housed in a 12-h light/dark cycle, with food and water ad libitum. Mice were euthanized and blood collected for serum preparation from 16 weeks old BL/6 animals, from 16 weeks old arthritic K/BxN animals and from 26 – 38 weeks old NZB/WF1 mice with established glomerulonephritis.

### Statistical analyses

Statistical analyses were performed using Graphpad Prism software (v6) and appropriate tests.

## Supplementary Materials

Fig. S1. rhTNFα is not toxic to mouse lymphocyte BW5147 cells even at high concentrations.

Fig. S2. FcγRs are activated by VEGFA and IL-5 sICs

Fig. S3. Distinct activation patterns of NK cells incubated with inverse sICs

Fig. S4. AF4 elution profiles of Ifx/TNFα-immune complexes.

Table S1. Analysis of the molar mass distribution of ICs from AF4 data.

## Acknowledgements

We thank T. Schleyer (IR-B Biobank) for providing patient samples. We are indebted to Falk Nimmerjahn (Universität Erlangen-Nürnberg) for providing KRNtg mice.

## Data and materials availability

All data associated with this study are present in the paper or Supplementary Materials.

## Author contribution

Conceived and designed the experiments: P.K., H.C., A.M-P., M.H.

Performed the experiments: P.K., H.C., M.H., A.M-P, U.S.

Analyzed the data: P.K., H.C., A.M-P, U.S.

Contributed/reagent/sample material: RE.V.

Writing and original draft preparation: P.K., H.C.

Review and editing: H.H., RE.V., P.K.

## Supplementary material

**Fig. S1.**
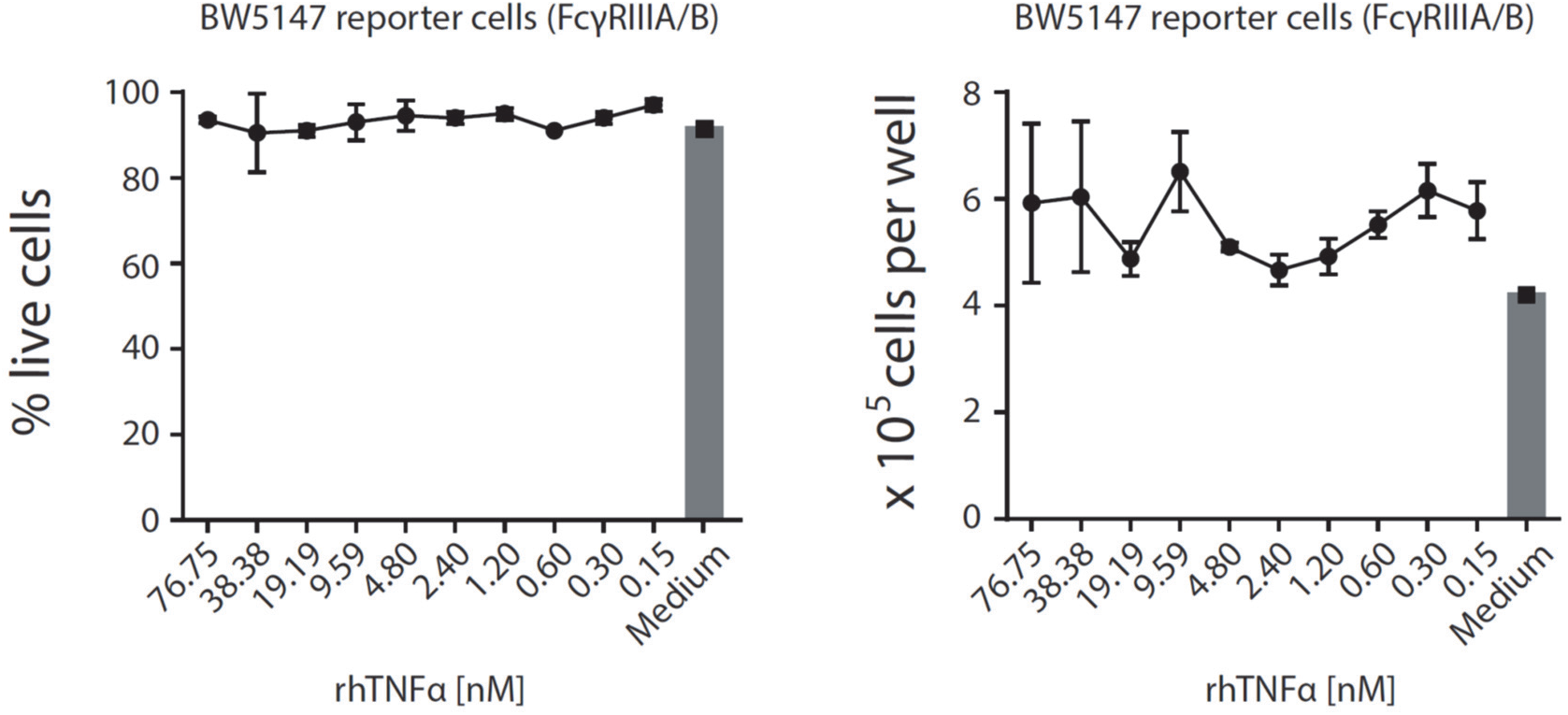
rhTNFα is not toxic to mouse lymphocyte BW5147 cells even at high concentrations. Cell count and percentage of live cells were unaltered over a 16 h time frame of reporter cell culture in the presence of indicated rhTNFα concentrations and comparable to regular growth in complete medium. Experiments were conducted in 3 replicates. Error bars = SD.

**Fig. S2.**
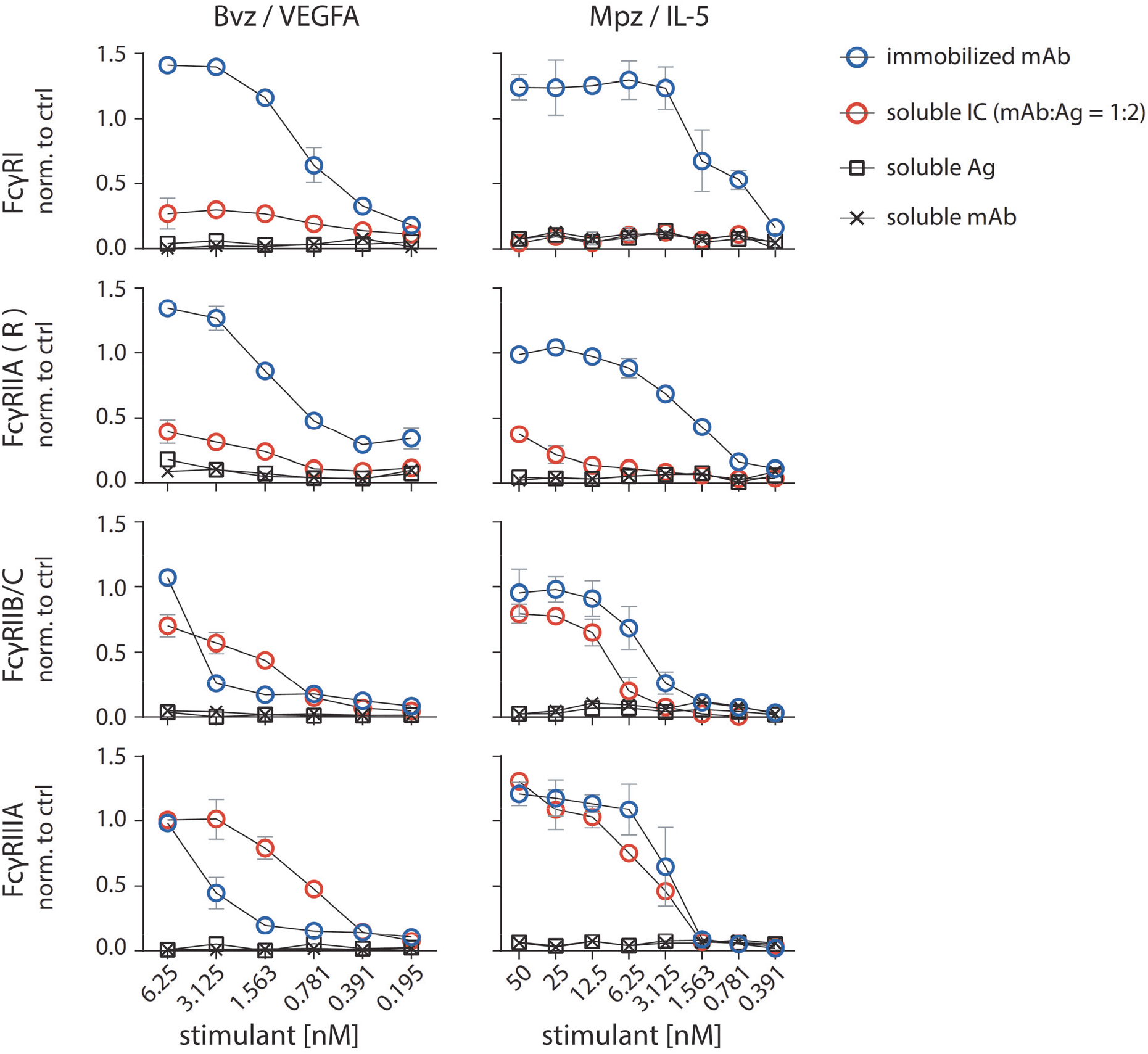
FcγRs are activated by sICs formed from multivalent antigens. Two different multivalent ultra-pure antigens (Ag) mixed with respective therapy-grade mAbs were used to generate sICs as indicated for each set of graphs (top to bottom). IC pairs: mepolizumab (Mpz) and rhIL-5; bevacizumab (Bvz) and rhVEGFA. X-Axis: concentrations of stimulant expressed as molarity of either mAb or Ag monomer and IC (expressed as mAb molarity) at a mAb:Ag ratio of 1:2. Soluble antigen or soluble antibody alone served as negative controls and were not sufficient to activate human FcγRs. FcγR responses were normalized to immobilized rituximab (Rtx) at 1 µg/well (set to 1) and a medium control (set to 0). Two independent experiments performed in technical duplicates. Error bars = SD. Error bars smaller than symbols are not shown.

**Fig. S3.**
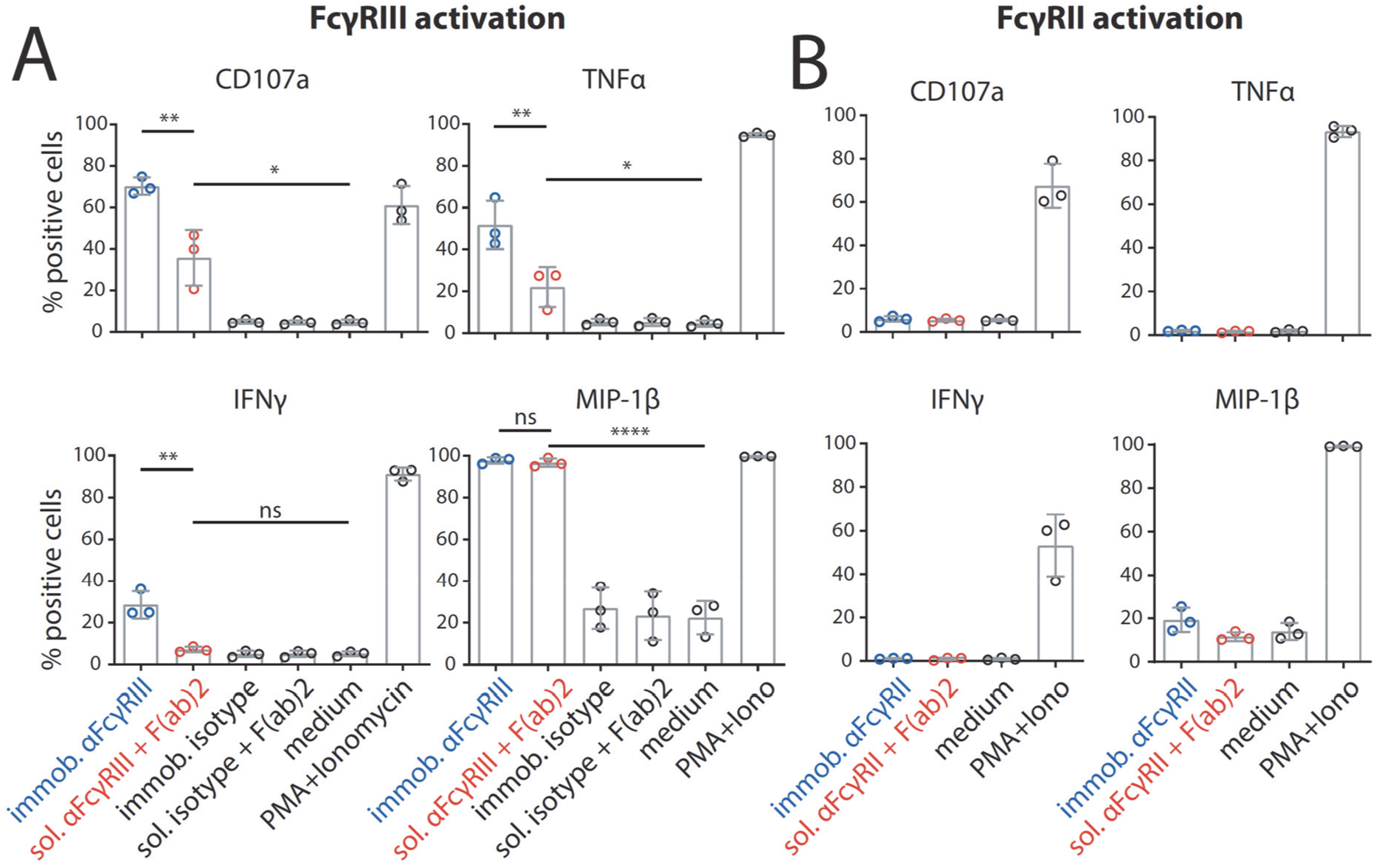
Distinct activation patterns of NK cells incubated with inverse sICs. Negatively selected primary NK cells purified from PBMCs of three healthy donors were tested for NK cell activation markers. Error bars = SD. One-way ANOVA (Tukey); *p<0.05, **p<0.01, ***p<0.001, ****p<0.0001. A) NK cells were incubated for 4 h with immobilized FcγRIII-specific mAb, soluble mouse-anti-human IgG F(ab)_2_ complexed FcγRIII-specific mAb (reverse sICs), immobilized IgG of non-FcγRIII-specificity (isotype control) or soluble F(ab)_2_ complexed isotype control (all at 1µg, 10^6^ cells). Incubation with PMA and Ionomycin served as a positive control. Incubation with medium alone served as a negative control. B) As in A using an FcγRII-specific mAb. NK cells from the tested donors in this study do not react to FcγRII activation.

**Fig. S4.**
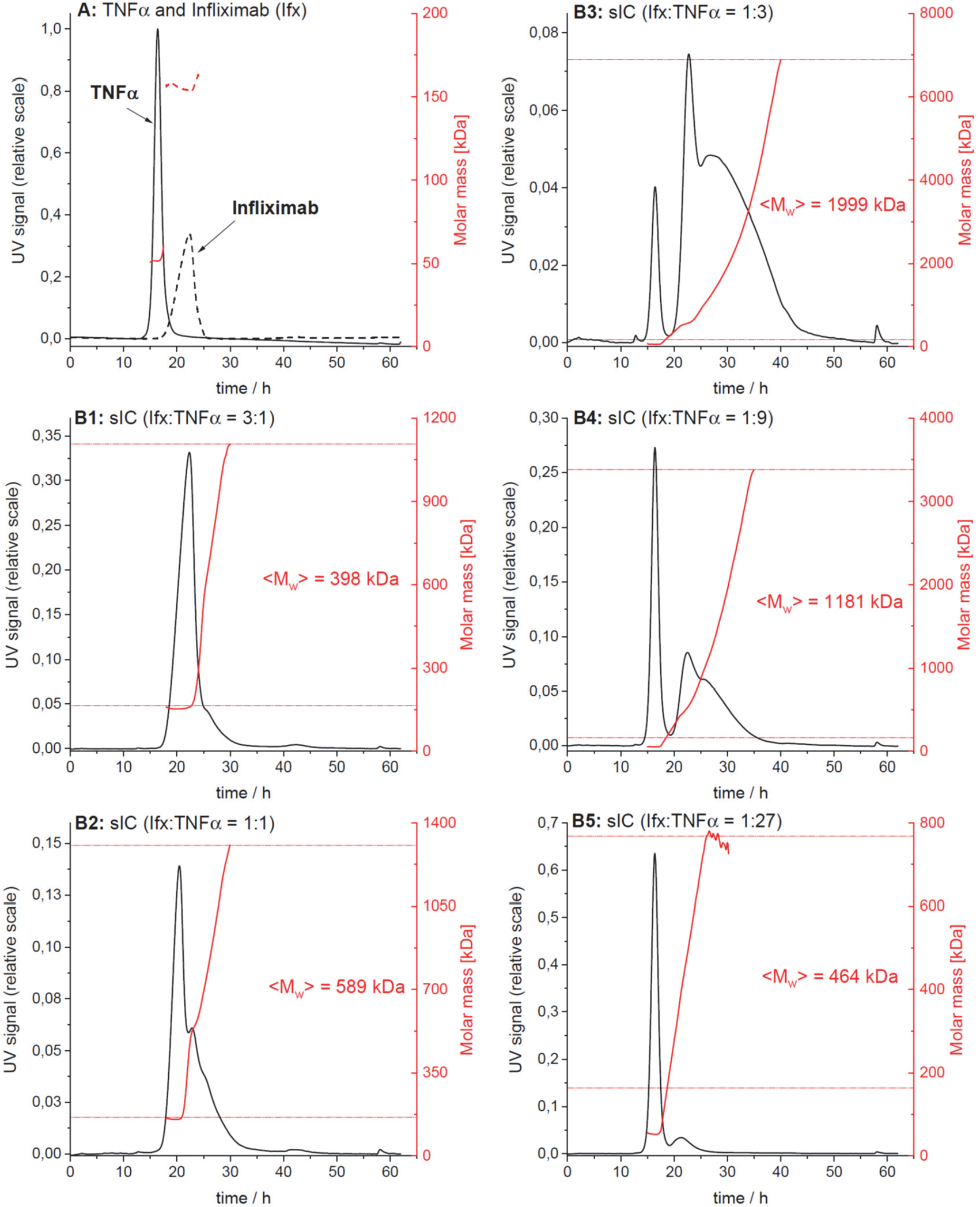
AF4 elution profiles of Ifx/TNFα-immune complexes. The elution profiles from one of three independent runs are shown. Protein concentration in the eluate is shown in black (UV signal at λ = 280 nm, normalized to the highest UV signal found in this experiment), molar masses determined by MALS for a given retention time in red. Horizontal red lines indicate the range of molar masses used to calculate the mass-weighted mean of molar masses <M_w_>. A) Overlay of the elution profiles obtained for TNFα and Ifx, respectively; B1 to B5) Elution profiles for sICs formed after incubation of TNFα and Ifx at different molar ratios.

**Table S1.** Analysis of the molar mass distribution of ICs from AF4 data. For a given elution time, the AF4 profiles provide the concentration (UV) at which a given molar mass (MALS) of a protein is present in the sample. The molar mass distribution of Ifx, TNFα and their immune complexes (sICs) was obtained by plotting the cumulative frequency as a function of molar mass. For a selected range of molar masses, a mass-weighted mean value (<M_w_>) was calculated. All detected molar masses were selected in the case of Ifx and TNFα whereas only molar masses larger than the maximal molar mass found for Ifx were assigned to sICs. The table shows the range of assigned molar masses and the calculated <M_w_> for each AF4 run (n = 3).

| Sample | Range of assigned molar masses [kDa] |  |  | Mass-weighted mean of assigned molar masses [kDa] |  |  |  |
| --- | --- | --- | --- | --- | --- | --- | --- |
| | Run 1 | Run 2 | Run 3 | Run 1 | Run 2 | Run 3 | Mean ± SD |
| Infliximab, IFX | 158 – 182 | 153 – 164 | 159 – 193 | 162 | 156 | 163 | 160 ± 4 |
| TNF -alpha | 52 – 55 | 51 – 61 | 52 – 62 | 52 | 52 | 52 | 52 ± 0 |
| Immune complexes |  |  |  |  |  |  |  |
| IFX/TNF 3:1 | 182 – 1.16$\cdot 10^3$ | 164 – 1.11$\cdot 10^3$ | 193 – 1.10$\cdot 10^3$ | 409 | 398 | 518 | 442 ± 66 |
| IFX/TNF 1:1 | 182 – 2.06$\cdot 10^3$ | 164 – 1.31$\cdot 10^3$ | 193 – 1.42$\cdot 10^3$ | 801 | 589 | 681 | 690 ± 106 |
| IFX/TNF 1:3 | 182 – 5.05$\cdot 10^3$ | 164 – 6.89$\cdot 10^3$ | 193 – 10.8$\cdot 10^3$ | 1.77$\cdot 10^3$ | 2.00$\cdot 10^3$ | 2.61$\cdot 10^3$ | 2.13$\cdot 10^3 \pm 435$ |
| IFX/TNF 1:9 | 182 – 5.36$\cdot 10^3$ | 164 – 3.38$\cdot 10^3$ | 193 – 3.51$\cdot 10^3$ | 1.66$\cdot 10^3$ | 1.18$\cdot 10^3$ | 1.17$\cdot 10^3$ | 1.34$\cdot 10^3 \pm 279$ |
| IFX/TNF 1:27 | 182 – 1.68$\cdot 10^3$ | 164 – 768 | 193 – 1.01$\cdot 10^3$ | 689 | 464 | 521 | 558 ± 117 |

